# USHER: Guiding Foundation Model Representations through Distribution Shifts

**DOI:** 10.1101/2025.11.20.689462

**Authors:** Aditya Pratapa, Purushothama Rao Tata, Rohit Singh

## Abstract

Foundation models pre-trained on certain biological data modalities exhibit systematic representational biases when encountering out-of-distribution (OOD) data from new assays. The embedding drift largely arises from instrumentation and protocol-related artifacts rather than true biological variation in cell states or tissue morphology. These drifts are distinct from conventional batch effects and cannot be remedied by retraining as sample sizes are often insufficient, and modifying existing embeddings breaks downstream tools that depend on stable representations. We introduce USHER, an adaptable framework to learn simple transforms that return OOD embeddings to a foundation model’s reference space. USHER enables embedding transformation via an expectation maximization-style procedure. Given a reference in-distribution sample, USHER first estimates a Fused Gromov-Wasserstein coupling that aligns unpaired OOD (source) and reference (target) embeddings by minimizing transport distance while preserving local structure. To make optimal transport couplings more useful for down-stream tasks, we introduce the concept of entropic filtering to retain only high-confidence correspondences. In the second step, USHER learns a low-complexity transformation that reliably restores the model’s representation space for OOD data. We demonstrate this learned transformation generalizes to other OOD data from similar experimental conditions. We applied USHER to correct platform-specific biases seen when running scGPT on Xenium transcript counts: USHER maps Xenium embeddings back to the native scRNA-seq representation space, improving cell type clustering and cross-platform integration. Histopathology foundation models trained on H&E images fail on MALDI metabolite-profiled tissue images due to data-acquisition artifacts. USHER corrects these, enabling cell-type classification and protein abundance imputation. USHER offers a generalizable framework to make biological foundation models portable across a rapidly-evolving experimental landscape.

## 1 Introduction

Foundation models (FMs) trained on massive biological datasets have demonstrated remarkable success in encoding cellular and tissue states [1,2,3]. We focus on FMs where input data originates from protocols and assays with diverse data distributions; these include single-cell transcriptomics FMs, chromatin accessibility models, and histopathology imaging FMs. We investigate two questions: how do these models perform when confronted with data from new protocols they were not pre-trained on? And if there are resulting challenges, can they be surmounted effectively? This is an important direction of research because new assays continue to be invented and FMs that can not extend to these would have limited usefulness.

Our key first contribution is demonstrating that the two foundation models we investigated—the single-cell model scGPT [4] and the H&E imaging model H-optimus-1 [5]—face substantial representational challenges with out-of-distribution (OOD) data. When transcript counts from Xenium spatial-transcriptomics assays are input to scGPT, the resulting embeddings differ from those obtained from single-cell RNA-seq (scRNA-seq) on the same tissue. Similarly, when H&E imaging follows MALDI mass spectrometry on tissue, the matrix crystallization and laser etching alter the image so that foundation model embeddings are distinct from standard H&E embeddings of the same tissue.

Notably, with OOD data, the resulting FM representations are *degraded*, not destroyed. A UMAP of scGPT embeddings of Xenium transcript counts does capture distinct cell types. However, when a single UMAP is fitted on data that contains both Xenium embeddings and scRNA-seq embeddings, the two sources separate. Given that a major focus of single-cell FM design has been addressing data-distribution variability from different sources (e.g., Geneformer [6] maps transcript counts to gene rankings, while scGPT [4] groups them into 50 bins), it appears that the models are able to discern the biological structure substantially, even if not entirely. We found a similar observation to hold for post-MALDI imaging data, as described later.

Our second key contribution is to show that conventional batch-effect correction and normalization methods are insufficient for this challenge. For single-cell data, we evaluated diverse scRNA-seq batch correction methods (Harmony, Scanorama, sysVI, ComBat), as well as a dedicated spatial-to-scRNA-seq mapper (Tangram). For H&E images, we applied Vahadane color normalization [7]. All methods failed to align the embeddings. We suspect this is for two reasons. First, embedding alignment is a strictly stronger requirement than matching phenotypes such as cell-type labels; the latter can be inferred from the former but not vice versa. Second, batch correction methods assume batches share a common data-generation process with only technical noise varying, allowing correction through centering, scaling, or non-linear warping. However, Xenium and standard scRNA-seq represent fundamentally different transcript-count distributions: Xenium covers far fewer genes (300 vs whole-genome) and its per-cell counts follow a different distribution than scRNA-seq. Even though the FM can retrieve the internal covariance structure of OOD data, it cannot map them to its original space.

We emphasize that this inability to obtain consistent embeddings for OOD data is a critical limitation. As exemplified by protein language models, many compelling applications of biological foundation models leverage their embeddings extensively.

In natural language processing and computer vision, distributional shifts are typically addressed through domain adaptation and model fine-tuning [8,9,10]. This is not an appealing approach for biological FMs. We typically have limited samples from new protocols, making fine-tuning difficult. There is also the risk of overfitting and catastrophic forgetting with such retraining [11]. More critically, fine-tuning risks altering the underlying embedding space, breaking compatibility with downstream tools and models built on the original embeddings. Instead, we propose to post hoc transform OOD embeddings back to the reference in-distribution space. We hypothesize this is feasible because FMs already capture substantial biological structure from OOD data; they simply map it to a different region of embedding space.

The third and most important contribution of this work is USHER (Unified Strategy for sHifting towards Established Representations), a general framework for post hoc transformation of foundation model embeddings from OOD data to the original reference space. Given a reference in-distribution dataset from similar biological context, USHER learns a transformation that aligns OOD embeddings with the reference distribution. The reference sample need not be identical to the OOD sample—we require only that similar biological states exist in both datasets. Importantly, the learned transformation generalizes: it applies not just to the specific sample used for training but also to other samples from similar experimental conditions.

USHER operates through an expectation-maximization framework that alternates between two steps until convergence. First, it establishes soft *correspondences* between OOD and reference embeddings using Fused Gromov-Wasserstein (FGW) optimal transport (OT), which aligns distributions while preserving their internal geometric structure. The precise FGW formulation is modality-specific. To make OT-based correspondence imputation more robust in high-dimensional biological datasets, we introduce *entropic filtering* to prune low-confidence matches, substantially improving robustness when source and target samples match only approximately. Second, it learns a low-complexity *transformation* (linear or nearly-linear) that maps OOD embeddings toward their matched references. The low-complexity constraint, with regularization towards the identity transformation, ensures interpretability and guards against overfitting when correspondence information is limited.

We demonstrate that USHER works effectively in two very distinct settings: single-cell transcriptomics and H&E histopathology foundation models. For single-cell data, we correct scGPT embeddings from Xenium spatial transcriptomics to align with standard scRNA-seq representation space. Using lung tissue with matched Xenium and scRNA-seq measurements, USHER achieves substantial improvement in cross-platform integration metrics compared to uncorrected embeddings and standard batch-correction or mapping methods and unlocks imputation of unassayed genes. For histopathology, we address post-MALDI H&E images where mass spectrometry introduces crystallization artifacts and laser damage. USHER learns a transformation to proteomics-linked H&E image that correct these artifacts in embedding space, recovering tissue classification accuracy and enabling protein abundance prediction from morphology.

## 2 The USHER algorithm

### Sketch of the algorithm

Given a FM trained on standard data *T* (e.g., scRNA-seq atlas) and embeddings from a new protocol *S* (e.g., Xenium spatial transcriptomics), USHER learns a transformation that maps *S*’s embeddings back to the FM’s native representation space (Fig. 1a). Because *S* and *T* are *unpaired*, we cannot use individual matches directly to learn the transform. Instead, USHER uses an expectation-maximization–style procedure with two alternating steps: estimating correspondences by optimal transport and fitting a low-complexity transform.

**Fig. 1:**
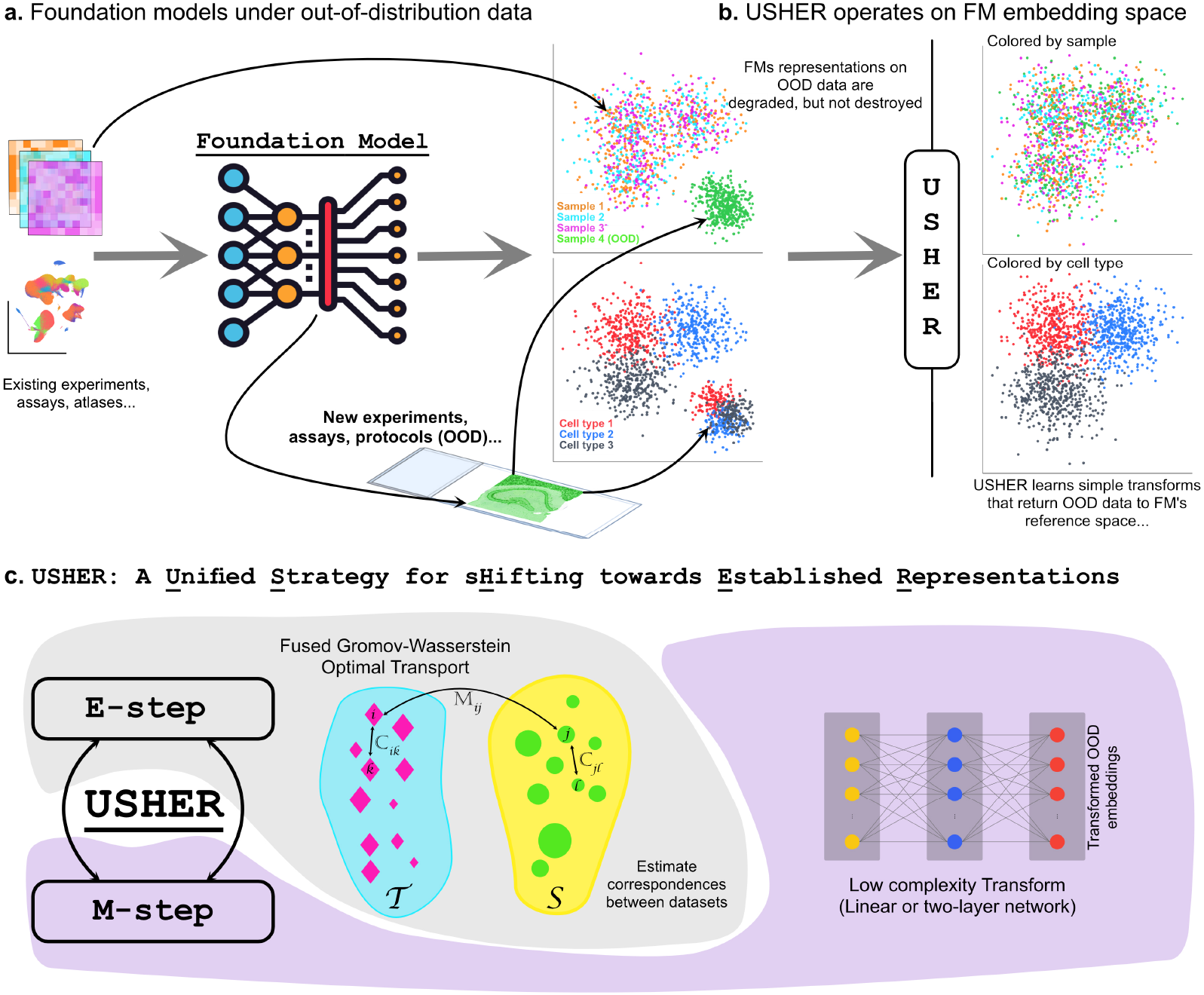
USHER Overview. **a** Foundation-model embeddings degrade under out-of-distribution (OOD) assays, experiments, and protocols. **b** USHER operates purely in embedding space, learning low complexity (e.g., linear) transforms that return OOD representations to the model’s established reference space. **c** USHER alternates between estimating cross-dataset correspondences via fused Gromov–Wasserstein optimal transport (E-step) and fitting a simple neural transform (M-step), yielding transformed embeddings for OOD data.

#### E-step (Correspondence via OT)

We compute soft correspondences *P* between source and target samples using Fused Gromov–Wasserstein optimal transport [12], which considers a combination of the within-domain structure and cross-domain similarity. In high-dimensional, partially-mismatched biological datasets, many OT couplings are diffuse and uninformative. We therefore introduce *entropic filtering*: pruning rows of *P* with high entropy to retain only high-confidence matches, then extracting a sparse mapping via Hungarian assignment. Unlike standard OT workflows that preserve soft couplings, this hardening step produces a sparse and interpretable correspondence set and empirically improves optimization stability (Fig. 1b). We believe entropic filtering could be broadly useful in the many settings where OT-based probabilistic couplings form the basis of a downstream task.

#### M-step (Transform Learning)

Given the filtered correspondences, we learn a parametric transformation *f*_*θ*_ that maps each OOD embedding to its matched reference. We implement *f*_*θ*_ as a shallow neural network (1 or 2 linear layers) with regularization toward the identity transform. This low-complexity design reduces the risk of overfitting when correspondence information is limited and induces only a modest recalibration of the original embedding rather than a wholesale re-embedding. When *f*_*θ*_ is strictly linear, it also becomes directly interpretable: as we show for scGPT, the learned transform is sparse and nearly diagonal. After each M-step update, we recompute cross-domain similarities using the transformed embeddings and iterate. The final *f*_*θ*_ acts as a reusable calibration that can generalize to new OOD samples from similar experiments without retraining (Fig. 1b).

### Problem Formulation and Notation

Let 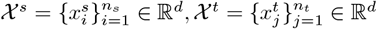denote the source (OOD) and target (ID) embeddings extracted from a frozen foundation model. Each 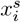 and 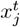 is a *d*-dimensional feature vector, and *n*_*s*_, *n*_*t*_ are the respective sample sizes. Crucially, no paired labels are available between *χ*^*s*^ and *χ*^*t*^; we only observe two independent samples from related but distinct distributions. Our goal is to learn a parametric mapping

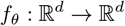

such that the transformed source distribution *f*_*θ*_(*χ*^*s*^) becomes geometrically and semantically aligned with the target distribution *χ*^*t*^ while keeping the foundation model fixed. Because explicit pairs are unavailable, we introduce a soft correspondence matrix *P* ∈ *Π*, where *Π* denotes the set of all *n*_*s*_ × *n*_*t*_ row-stochastic matrices. The entry *P*_*ij*_ represents the degree to which source point 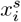 is associated with target point 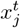.We formulate this as a joint optimization *f*_*θ*_ and *P*:

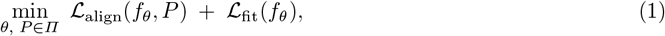

which we solve using an EM-style procedure: the E-step updates *P* via optimal transport, and the M-step updates *f*_*θ*_ given the current correspondences.

### Expectation Step: Optimal Transport–based Correspondences

Given the current transformation *f*_*θ*_ (initialized as the indentity map), we estimate a soft correspondence matrix *P* between the transformed source embeddings *f*_*θ*_(*χ*^*s*^) and the target embeddings *χ*^*t*^. To do this, USHER uses *Fused Gromov–Wasserstein* (FGW), which combines two complementary notions of similarity:

- Within-domain structure (ℂ^*s*^, ℂ^*t*^) is captured by comparing the intra-domain distance patterns within *χ*^*s*^ and *χ*^*t*^. Following SCOTv2 [13], we compute pairwise structure using *k*NN-graph (*k* = 30) geodesic distances [13], where edges are formed using cosine similarity of the FM embeddings. This yields structure matrices ℂ^*s*^ and ℂ^*t*^ with entries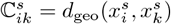 and 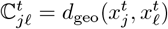where *d*_geo_ is the shortest–path distance on the *k*NN graph. These graph-geodesic distances provide a modality-agnostic notion of local geometry that remains comparable across domains even when absolute embedding locations are shifted.
- Cross-domain similarity (*M*) between individual points, encoded in a combined cost matrix. We first define a feature cost

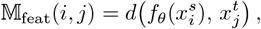

where *d* is typically cosine or Euclidean distance. Because the source and target embeddings generally lie in shifted or non-comparable regions of the foundation model space, USHER also constructs an auxiliary cost *M*_aux_(*i, j*) from features that live in a shared domain (such as estimated cell-type probabilities or spatial coordinates, Sec. A.1). The two are combined as

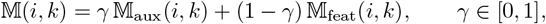

where larger *γ* emphasizes the auxiliary structure and smaller values place more weight on the foundation-model embeddings. At the first iteration, we warm–start the alignment using only the auxiliary cost, following standard practice in cross-domain transport methods to avoid circularity before the two embedding spaces have a shared reference frame.

We combine the feature and structural terms in a Fused Gromov-Wasserstein (FGW) objective:

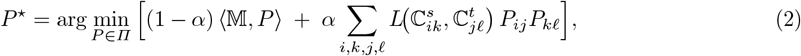

where *α*∈ [0, 1] interpolates between pure feature OT and pure Gromov–Wasserstein (GW): *α* = 0 recovers just feature-based OT, whereas *α* = 1 yields pure GW, which matches points based solely on structural geometry defined by the structure matrices ℂ^*s*^ and ℂ^*t*^. *L*(·, ·) is a point-wise loss (we use the squared loss). As is standard practice, we solve an entropically regularized (*ϵ* = 0.05) version of eq. (2) to obtain numerically stable, Sinkhorn-style GPU solvers for large problem sizes.

FGW solves for the correspondence matrix *P* that best balances these feature-level and structure-level agreements under the chosen (*α, γ*). Because the alignment problem is global rather than partial, we use the balanced FGW formulation, which avoids discarding rare states and ensures that the entire source distribution participates in the correspondence. If necessary, we sub-sample one distribution.

After computing the soft correspondence matrix *P*, we apply an entropic filtering step (as described previously) that removes high-entropy, low-confidence rows (typically the lowest ∼10% by entropy). On the remaining subset, we solve a linear sum assignment (Hungarian) problem on *P* to obtain a sparse one-to-one set of high-confidence pairs, which we use to drive the subsequent M-step update of *f*_*θ*_.

### Maximization Step: Learning a Low-Complexity Transform

Given the filtered correspondence matrix *P* and the one-to-one matches extracted by the Hungarian step, the M-step updates the parameters of *f*_*θ*_ so that transformed source embeddings move toward their matched targets. We parameterize *f*_*θ*_ as a low-complexity feedforward mapping (typically a single linear layer or a shallow one-hidden-layer MLP). This keeps the number of parameters small relative to *d* and empirically avoids wholesale distortion of the foundation-model embedding geometry.

Let *π*(*i*) denote the target point matched to source point *i* by the Hungarian assignment applied to the filtered correspondence matrix *P*. The M-step minimizes a fitting loss of the form

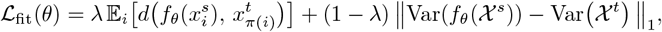

where *d* is Euclidean distance. The first term encourages each source embedding to move toward its matched target under the learned transformation, while the variance term keeps the overall feature-wise dispersion of *f*_*θ*_(*χ*^*s*^) close to that of *χ*^*t*^, preventing mode collapse.

We update *θ* using Adam for a small number of gradient steps per EM iteration. In practice, this low-capacity parameterization and variance regularization yield modest, stable shifts that correct domain drift while preserving the semantic structure encoded by the foundation model.

### Implementation details, Hyperparameters and Runtime

We standardize all embeddings using target-domain statistics before computing distances or training, then invert this standardization after transformation to maintain original units (Sec. A.3). Since USHER only needs to learn the global transform *f*_*θ*_ rather than dense correspondences for all points, we run FGW on representative subsamples selected via geometric sketching [14] for single-cell data or spatial windowing for imaging data (Sec. A.2).

USHER requires three key hyperparameters. The FGW trade-off *α* balances feature similarity against structural preservation, with values between 0.3–0.7 performing well across experiments. The auxiliary weight *γ* controls the relative importance of domain-invariant signals (cell types or spatial coordinates) versus foundation model embeddings in the cost matrix *M* = *γM*_aux_ + (1 − *γ*)*M*_feat_. The fitting loss weight λ determines whether the M-step prioritizes pointwise alignment of matched pairs or matching overall feature dispersion between *f*_*θ*_(*χ* ^*s*^) and*χ* ^*t*^. We found USHER robust to these parameters across reasonable ranges (Sec. A.4). Other settings remain fixed: sketches of ∼ 4, 000 points per sample, 100 M-step epochs per EM iteration with Adam optimizer (learning rate 10^*−*3^), and 25 total EM iterations. The method runs efficiently on a single GPU, processing ∼30,000 cells under 5 minutes per EM iteration on a single Nvidia A6000 GPU.

## 3 Results

We evaluated USHER on two foundation models. First, we applied it to scGPT embeddings of Xenium spatial transcriptomics data. Typical Xenium gene panels are small (∼300–500 genes) and the ability to map their FM embeddings to a reference scRNA-seq space would enable, among other applications, gene and program imputation. Second, we applied USHER to H&E histopathology images. In spatial omics, H&E often serves as an auxiliary readout available with Visium, PhenoCycler-Fusion (PCF), and MALDI, providing a way to align independent measurements. However, techniques like MALDI introduce crystallization and laser etching artifacts that degrade H&E image embeddings, creating an integration challenge.

### 3.1 Applying USHER to adapt scGPT embeddings of Xenium transcripts

We used matched human idiopathic pulmonary fibrosis (IPF) lung datasets from Vannan *et al*. [15], comprising a Xenium section (38,280 cells profiling 343 genes) and a dissociated scRNA-seq atlas from matched tissue. We obtained 11 broad cell classes for the scRNA-seq reference based on the original study’s expert-curated annotations [15]. To assess initial alignment quality and initialize our EM algorithm, we transferred these labels to Xenium with a logistic-regression classifier trained on genes common to both platforms. We neither need nor expect this label-transfer to be very precise; it only needs to be approximate. To mitigate mass imbalance during optimal transport, we stratified the scRNA-seq cohort to match Xenium’s marginal cell-type distribution, yielding 26,323 reference cells for analysis (Sec. A.5).

For each assay separately, the scGPT embeddings [4] capture biologically meaningful structure (Fig. 2a– d), with major alveolar, immune, and stromal populations clearly identifiable. Xenium clusters did exhibit less separation and greater local dispersion (Fig. 2b) than scRNA-seq clusters (Fig. 2a), likely reflecting the lower per-cell transcript detection in Xenium 2(e). But when visualized jointly, the distributional shift is clear, with the two datasets occupying distinct regions of the embedding space (Fig. 2c–d).

**Fig. 2:**
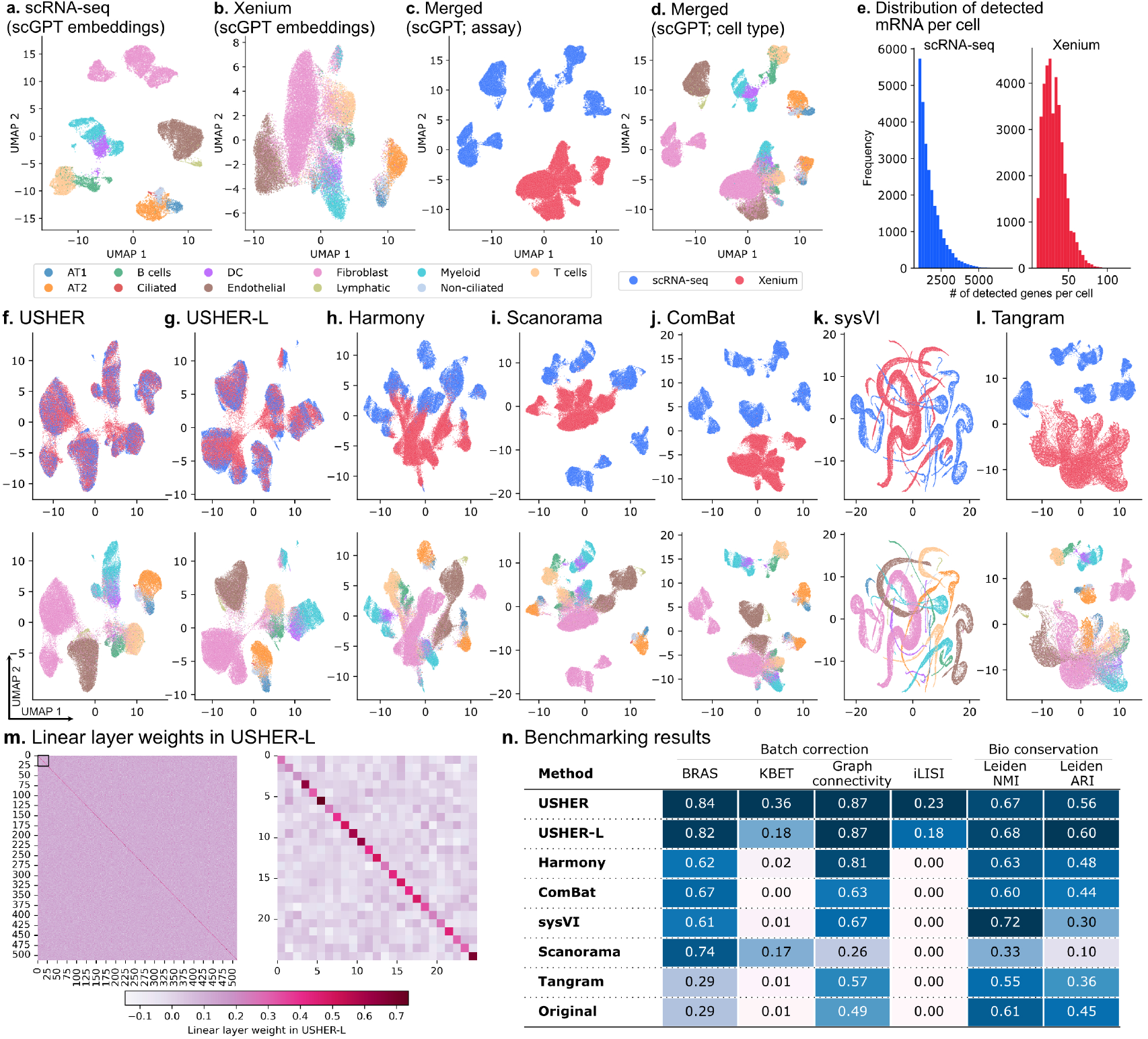
Assay mixing performance across integration methods. UMAPs of **a–d** show the original scGPT embeddings. **e** Distribution of mRNA counts per cell. Panels **f–l** show UMAPs after integration with USHER, USHER-L, Harmony, Scanorama, ComBar, sysVI, and Tangram, colored by batch (top) and cell type (bottom) for scRNA-seq and Xenium datasets. **m** USHER-L’s learned W matrix for the 512-dimensional scGPT embeddings (left) and a zoom-in of the top-left 25× 25 block (right), highlighting its strong diagonal structure. **n** Assay mixing performance across multiple metrics.

#### Batch-effect correction (BEC) methods fail to resolve the discrepancy

We assessed BEC and related methods to harmonize these embeddings, choosing four popular methods with very distinct approaches: Harmony [16], Scanorama [17], ComBat [18], and sysVI [19]. We also evaluated Tangram [20], a dedicated spatial-to-scRNA-seq mapper. To compute a comprehensive array of metrics, we used the ‘scib-metrics’ package [21,22], which provides standardized integration benchmarks with all metrics normalized so that higher values indicate better performance. The package computes a *batch correction score* (aggregating metrics like batch ASW, graph connectivity, and iLISI) and a *bio conservation score* (aggregating metrics like cell-type ASW, NMI, and ARI). In particular, scIB authors recommend the BRAS metric (batch-removal-adapted silhouette width) as a good balance of batch-mixing with biological structure preservation [22].

While all tested methods did reduce the separation between Xenium and scRNA-seq embeddings, they failed to achieve meaningful integration (Fig. 2h–l; Fig. 2n). Scanorama achieved the closest alignment to USHER in terms of BRAS score but its post-integration modalities still remained largely separate in UMAP space (iLISI=0) (Fig. 2c). ComBat, sysVI and Harmony performed comparably or worse. Tangram, despite being specifically designed for spatial-to-scRNA-seq mapping, also failed to produce a suitable shared embedding space. One reason for these failures may be that we applied these methods to embedding space, which is necessary since our goal is to obtain matched embeddings. We also tried using Tangram to map transcripts directly and then compute scGPT embeddings, but this also fared poorly (Sec. A.6, Fig. S1). Moreover, Xenium and scRNA-seq represent fundamentally different transcript-count distributions: Xenium covers fewer genes and its per-spot counts are sparser (Fig. 2(e)). BEC methods, which mainly seek to address technical noise, seem to lack the machinery to bridge this OOD gap originating from scGPT’s black-box nature.

#### USHER effectively maps scGPT OOD embeddings to reference space

Applying USHER (Fig. 2f) realigns the Xenium embeddings into the reference scRNA-seq manifold while preserving the original biological clustering. We constructed the cross-sample cost matrix *M* using cosine similarity between scGPT embeddings; while initially approximate, these estimates become progressively more accurate over iterations of EM. We constructed the within-sample structure-preserving costs ℂ_*S*_ and ℂ_*T*_ using *k*-nearest neighbor graphs (*k* = 30). In the E-step, we set the FGW *α* = 0.3 (Fig. S2); in the M-step, λ = 0.9 (Fig. S3). To initialize correspondences (iter = 0), we introduce a cell-type matching based auxiliary term in *M*. In later iterations, we fix its weight at *γ* = 0.4 but ablations show that even *γ* = 0 after the initial iteration yields nearly identical performance (Fig. S4), i.e., the auxiliary signal is only needed to initialize, not to drive the alignment itself. USHER achieves a BRAS score of 0.84, higher than any of the methods under consideration. It also maintains strong NMI with unsupervised clustering (0.67) and strong graph connectivity and batch mixing (graph connectivity: 0.86, KBET: 0.36) (Fig. 2n; Fig. S5). Taken together, these metrics indicate that USHER successfully projects Xenium embeddings into the scRNA-seq reference space while preserving the biological structure captured by scGPT.

#### Ablations

To test robustness to the choice of mapping function, we constrained the transform to a single linear layer (**USHER-L**; Fig. 2g), replacing the two-layer neural network in the M-step. USHER-L achieved comparable performance to the full USHER model: BRAS of 0.82 versus 0.84, KBET of 0.36 vs 0.18, graph connectivity of 0.87 for both, and similar NMI (0.67 versus 0.68). The critical advantage of a linear layer is interpretability. Here, the USHER-L weight matrix turns out to be near-diagonal (Fig. 2m). This suggests that only subtle axis recalibrations were needed to adapt the OOD embeddings; the FM had already extracted the relevant biological signal.

USHER is robust to a range of hyperparameter choices of *α, γ*, and λ. Varying the FGW trade-off *α* shows that BRAS remains nearly unchanged for *α* ∈ [0.0, 0.9], while iLISI and KBET both peak at *α* = 0.3 (Fig. S2). Pushing *α* toward 1, i.e., reducing the objective to a pure GW structural match, leads to broadly degraded BRAS, iLISI, and KBET performance. Without the cross-sample term (*M*), the model aligns local geometries but fails to establish biologically meaningful correspondences. For the M-step correspondence weight λ, we observe a plateau of strong batch-mixing performance for λ ∈ [0.6, 0.9] (Fig. S3). Finally, the auxiliary-signal weight *γ* shows minimal effect on BRAS across its full range; iLISI and KBET are maximal at *γ* = 0.4, and performance remains stable for *γ* ∈ [0.0, 0.9] (Fig. S4). We also find that smaller values of *α* or *γ* increase the influence of feature-based matching (*M*_feat_) in the E-step, leading to more stable convergence across iterations (Fig. S6).

#### USHER mappings transfer to other samples from the same protocol

Since USHER seeks to learn the *distributional* rather than *sample-specific* characteristics of the new domain, we hypothesized that an USHER mapping fitted for one OOD sample would also be applicable to additional samples from the same experimental protocol. To test this, we applied the fitted USHER transformation—without retraining—to four additional Xenium sections from the same Vannan *et al*. cohort [15]. The resulting embeddings projected cleanly into the reference manifold, producing well-mixed UMAPs in which cells from all Xenium sections coherently overlapped with the scRNA-seq reference (Fig. 3a–b). Quantitatively, the held-out samples maintained high BRAS and KBET scores (Fig. 3d), indicating strong biological concordance and batch mixing across samples. One sample did show a lower iLISI value (0.04) though its BRAS and KBET scores still remained high. The aberrant iLISI score may be because the section had twice the number of cells and fixed-*k*NN metrics like iLISI are sensitive to this. Overall, these results indicate that USHER captures the systematic distributional shift introduced by this Xenium protocol, enabling the fitted transform to serve as a reusable calibration for future datasets with similar gene panel designs.

**Fig. 3:**
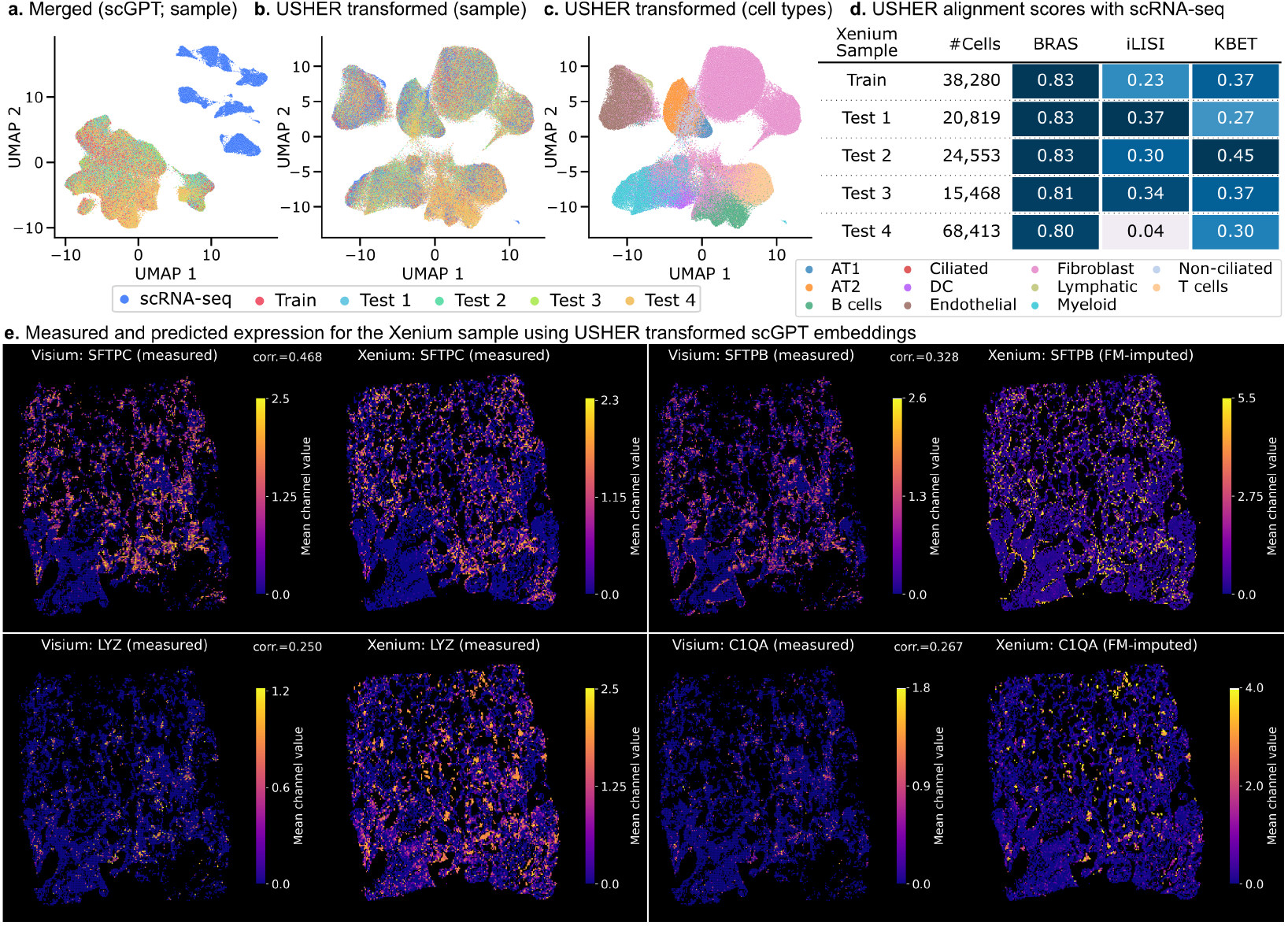
USHER generalizes to held-out samples and enables gene imputation. **a–c**. UMAP representation of USHER-transformed embeddings for held-out Xenium samples showing integration with scRNA-seq reference. **d**. Pairwise USHER alignment scores of Xenium and scRNA-seq samples. **e**. Spatial expression patterns for genes measured (left) in both Xenium and VisiumHD, showing baseline correlation. Imputed expression patterns for genes not in Xenium panel (right), showing comparable correlation to ground truth VisiumHD measurements.

#### USHER mappings enable gene imputation in Xenium data

Post-transformation, the standardized scGPT embedding enables cross-assay information transfer. As a demonstration, we applied USHER to impute expression of genes not in the original Xenium panel, assessing them against VisiumHD data also acquired on the same lung sample [15]. VisiumHD provides whole-transcriptome spatial profiling at near-cellular resolution, serving as ground truth. Notably, the baseline level of agreement between Xenium and VisiumHD is modest, reflecting assay variability: for two gene measured in both assays, we compared Xenium-measured and VisiumHD-measured expression for two genes measured in both assays, *SFTPC* and *LYZ*, the correlations were modest: *ρ* = 0.47 and *ρ* = 0.25, respectively. We next imputed expression for two genes not in the Xenium panel but measured in VisiumHD: *SFTPB* and *C1QA*. After transforming Xenium embeddings with USHER, we performed a 10-nearest-neighbor search in the scRNA-seq reference embedding space and assigned Xenium gene-expression estimates by averaging the expression profiles of the closest matching scRNA-seq cells. The imputed patterns are concordant with VisiumHD measurements, with Pearson correlations for SFTPB and C1QA (*ρ* = 0.33 and 0.27, respectively) comparable to the directly-measured genes (Fig. 3e–f). Visual inspection confirms that the imputed spatial localizations recapitulate known tissue structure: *SFTPB* marks alveolar type II cells, while *C1QA* localizes to macrophages.

### 3.2 Applying USHER to OOD histopathology data

To test USHER’s cross-modal generalizability, we applied it to histopathology images where the distributional shift stems from sample processing artifacts. We analyzed near-serial sections obtained from the lung adenocarcinoma tissue [23]. One section was H&E stained and imaged after MALDI mass spectrometry imaging (“post-MALDI”), while another was acquired from the same tissue block after PCF protein imaging (“post-PCF”). The post-PCF H&E represents the in-distribution case, processed using standard histological protocols. In contrast, post-MALDI has substantial artifacts: matrix crystallization from the organic matrix applied to the tissue surface, laser etching from the ionization process, and thicker sectioning required to withstand the MALDI procedure (Fig. 4a–d). These perturbations alter texture, color, and microstructure in ways that corrupt foundation model embeddings; e.g., note the laser etching’s cross-hatch pattern in Fig. 4d.

**Fig. 4:**
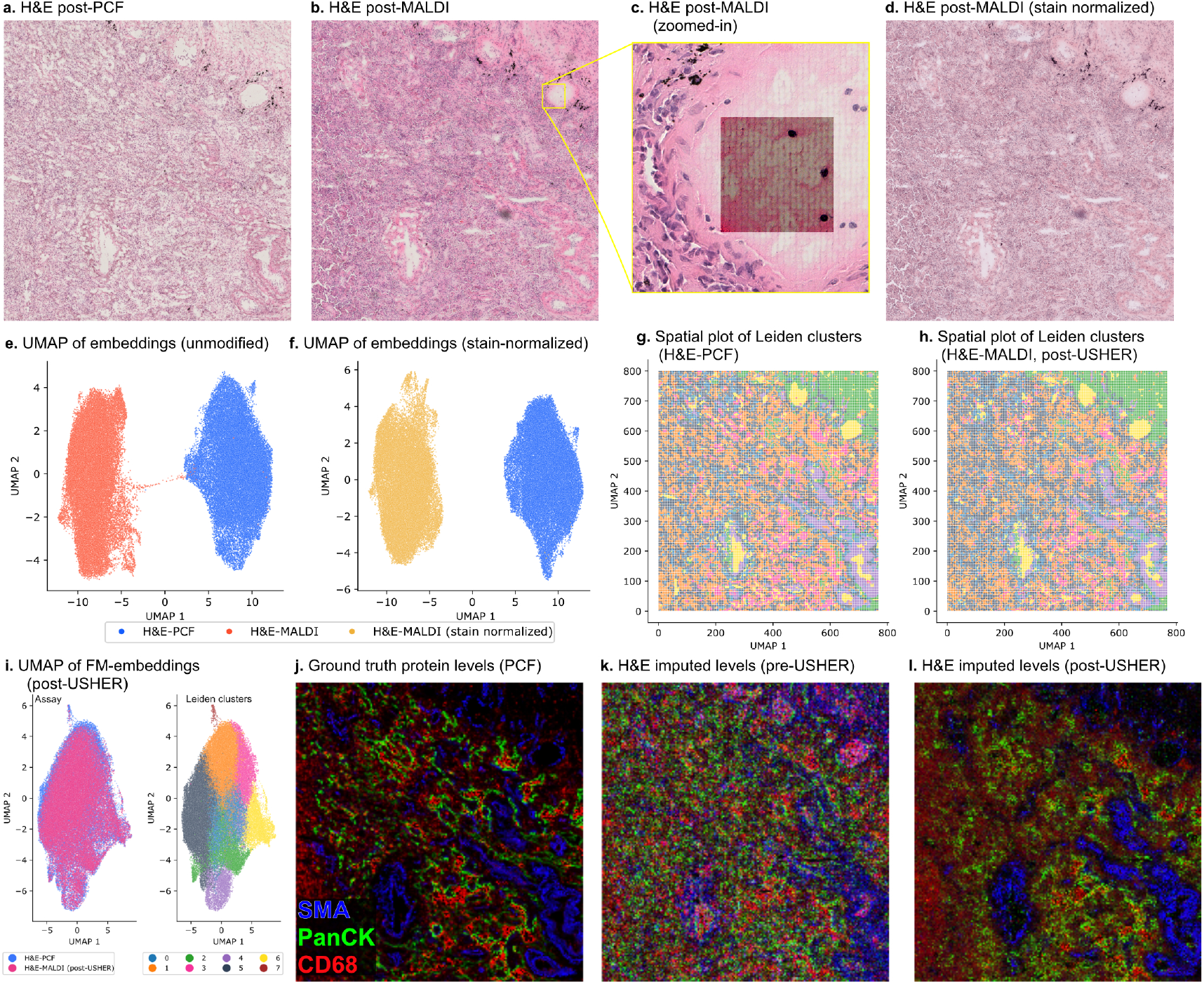
USHER corrects MALDI-induced artifacts in histopathology embeddings. **a–b**. H&E images for H&E acquired post-PCF (in-distribution, left) and H&E acquired post-MALDI (OOD, right) sections. **c**. Zoomed-in region of H&E post-MALDI showing MALDI-related artifacts. **d**. Vahadane color normalization of post-MALDI H&E image using H&E-PCF color space. **e–f**. UMAPs showing clear separation of FM-embeddings obtained on unmodified images (left) and Vahadane stain normalizated image (right). **g–h** Spatial clustering of corrected embeddings recovers morphologically coherent regions. **i**. USHER-corrected embeddings align post-MALDI to post-PCF manifold. (**j–l**) PCF protein abundance (left) and their prediction from raw embeddings (center) versus USHER-corrected embeddings (right) for SMA, PanCK, and CD68, showing recovery of spatial localization patterns.

When we extracted embeddings from both post-MALDI and post-PCF slides using H-optimus-1, the two embeddings occupied distinct regions of the representation space when visualized jointly in a UMAP. We first tested whether standard color normalization could resolve this separation, applying Vahadane stain normalization [7]. This method was specifically designed to correct for color variations in H&E images by separating hematoxylin and eosin stains and normalizing their distributions. Here, Vahadane normalization alone does not correct the embedding drift. The post-MALDI embeddings remain separated from post-PCF even after color normalization (Fig. 4e–f), indicating that the distributional shift is driven by physical artifacts rather than staining intensity.

#### USHER effectively maps H-optimus-1 OOD embeddings to reference space

We applied USHER to learn a transformation from post-MALDI to post-PCF embedding space. Details of the datasets, spatial batching strategy, as well as the auxiliary signal in *M* (it scores spatial proximity between source and target pixels), are in the Supplementary sections A.1–A.5. Applying USHER realigns the post-MALDI embeddings to the post-PCF manifold while preserving the fine-grained spatial organization of histological structures (Fig. 4g). Our PCA projection results (Fig. S7) further illustrate this effect: unmodified H&E-MALDI embeddings fail to reproduce the spatial structure captured in the PCF reference, whereas post-USHER embeddings recover the variation observed in H&E-PCF for PC-0. Assay mixing metrics like BRAS (0.91) and KBET (0.78) also showed improvement compared to original, unmodified embeddings (BRAS=0.68, KBET=0.0; Fig. S8). Similar to the scGPT case, constraining the USHER transform to be only linear (USHER-L) also degraded performance here (Fig. S8). Clustering of the USHER-corrected embeddings yields spatial patterns that closely match the reference PCF-based cell clusters (Fig. 4h–i), demonstrating recovery of morphologically coherent regions.

#### USHER enables protein abundance prediction from post-MALDI H&E

The ability to accurately estimate protein levels (and hence, cell types) from post-MALDI H&E is particularly valuable for spatial omics workflows, as it enables integration of mass spectrometry metabolite/lipid profiles with proteomic and transcriptomic readouts. We demonstrate how USHER-based embedding standardization conveniently enables such workflows. Here, the PCF assay provided spatially-resolved measurements of protein markers at subcellular resolution. We trained a regression model to predict protein marker intensities from H-optimus-1 embeddings of the *post-PCF* H&E data, then applied this model to predict protein levels from the USHER-corrected embeddings of *post-MALDI* H&E. We focused on three markers representing distinct tissue compartments: smooth muscle actin (SMA) marking smooth muscle and myofibroblasts, pan-cytokeratin (PanCK) marking epithelial cells, and CD68 marking macrophages (Fig. 4j). Predictions based on the raw, uncorrected post-MALDI embeddings failed to reproduce meaningful spatial localization: the predicted protein patterns showed little correspondence to the expected tissue structures (Fig. 4k). In contrast, predictions from USHER-transformed embeddings successfully recovered tissue boundaries and marker-specific spatial distributions (Fig. 4l), restoring morphological information that would otherwise be lost to MALDI-induced image degradation.

## 4 Discussion

Foundation models offer a scalable framework for integrating massive, diverse, and heterogenous biological datasets as demonstrated by the growing impact of protein and genomic language models [24,25,26,27,28]. However, single-cell and imaging foundation models face a unique challenge: protocol variations within a single modality far exceed those seen for protein/genomic language models. New experimental protocols will continue to emerge, creating distribution shifts that pre-trained models cannot anticipate.

Classical FM domain adaptation is undesirable because finetuning needs more data than new protocols typically provide, risks catastrophic forgetting, and can shift the embedding space that downstream tools depend on [10,11]. USHER overcomes these limitations by recognizing that the FMs do extract portions of the underlying biological structure in OOD data, but map it away from the reference embedding space. Consequently, adaptation then becomes a recalibration problem rather than relearning.

This insight drives our technical design. Low-complexity transforms reduce the risk of overfitting and remain interpretable. The resulting sparse, near-diagonal matrix suggests that only modest axis adjustments may suffice. And because paired in-distribution and OOD samples are rarely obtainable in practice, we employ optimal transport to infer correspondences, which naturally leads to our EM framework. Entropic filtering stabilizes this process by pruning uninformative couplings that would otherwise impair convergence and distort underlying biological structure. The fitted transform immediately enables new analyses: by mapping Xenium and post-MALDI H&E embeddings into respective reference spaces, USHER enabled imputation of unmeasured molecular features. This cross-protocol imputation capability will become increasingly critical as spatial omics technologies proliferate, each with unique measurement constraints.

A failure mode for USHER arises when the FM cannot extract even partial structure from OOD data. Protocols that fundamentally alter the measurement process might require preprocessing before USHER can be applied. Like batch correction methods, USHER carries a risk of overcorrection, potentially collapsing genuine biological variation while aligning distributions. However, USHER explicitly models assay-specific properties, it is therefore more likely to generalize across protocols rather than overfit to individual datasets. As more samples from a protocol accumulate, the M-step can fit across multiple examples, further improving generalizability.

USHER’s success across two fundamentally different modalities, transcriptomics and histopathology, suggests broad applicability. For new modalities and FMs, the framework offers a mix of tunability and stability: it accommodates modality-specific expertise in the OT formulation (E-step) while the M-step remains unchanged, and the approach appears robust to a range of hyperparameter choices, reducing the burden of protocol-specific tuning.

## Code and Data availability

The USHER code is publicly at https://github.com/rohitsinghlab/USHER.

## Acknowledgements

The authors gratefully acknowledge the support of the Chan Zuckerberg Initiative.

## A Supplementary Material

### A.1 Auxiliary Cross-Domain Cost *M*_aux_

To provide a stable, domain-invariant signal for initializing the correspondence map, USHER constructs an auxiliary representation for both domains and uses it to define an auxiliary cross-domain cost matrix *M*_aux_. For the two domains explored in this study (gene expression and spatial imaging), we provide principled procedures to construct this auxiliary signal when it is not already available

For single-cell assays, we learn a lightweight logistic–regression classifier on the scRNA-seq using genes shared with Xenium assay, and apply it to the Xenium sample to obtain predicted cell-type probability vectors. For spatial datasets, we first affine-align the near-serial tissue sections and use the resulting Euclidean distances between aligned spatial coordinates as auxiliary features. These constructions yield high-quality, low-dimensional quantities that capture shared biological or spatial structure even when the foundation-model embeddings are substantially shifted across domains. Notably, these auxiliary quantities are computed only for the samples used during training of *f*_*θ*_; once the transformation is learned for an assay, no auxiliary information is needed for inference.

Formally, let 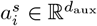 and 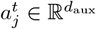 denote the auxiliary representations for source point *i* and target point *j*, respectively. We define the auxiliary cost as the pairwise Euclidean distance

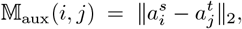

and normalize it for numerical stability:

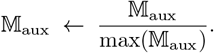

Incorporating *M*_aux_ complements the FGW feature and structure costs by encouraging correspondences that are not only structure-preserving but also similar in a shared, cross-domain signal. For spatial features, this biases the alignment toward physically proximal locations; for cell-type probability vectors, it favors phenotypically coherent matches. Following standard warm-start practice in cross-domain transport, USHER sets *γ* = 1 initially and learns the correspondences using *M*_aux_ alone. After the first iteration our ablations (Fig. S4) show setting *γ* = 0 and relying purely on *M*_feat_ still results in equally effective representations.

#### A.2 Batching Strategies

USHER requires pointwise correspondences to learn the *global* transform *f*_*θ*_, but these need not be computed densely for all cells. Instead, we estimate correspondences on representative subsets using two batching strategies that ensure each FGW solve operates on a compact yet structurally faithful portion of the data, enabling stable local correspondence estimation that jointly informs the global transform.

#### Geometric Sketching (Geosketching)

For scRNA-seq data, we use geometric sketching [14] to select diverse, representative subsamples that preserve the overall data structure:

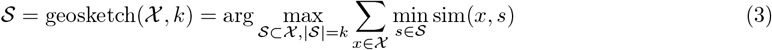

where *k* is the sketch size and sim(·,·) is a similarity measure. This iteratively selects points that maximize coverage of the feature space, ensuring each batch contains a representative sample of all cell types and states while maintaining computational efficiency. The sketching is performed independently on source and target to create matched batches for alignment.

#### Spatial Windowing

For spatial datasets, we partition the tissue into overlapping windows with overlap parameter, *δ* ∈ [0, 1] (typically 0.1). Windowing limits FGW to local neighborhoods where spatial structure varies smoothly, reducing ambiguity and computational cost. Correspondences are estimated within each window, and these local alignments collectively inform the global transform.

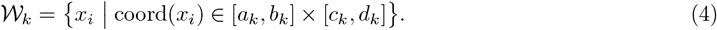

Here, [*a*_*k*_, *b*_*k*_] × [*c*_*k*_, *d*_*k*_] denotes the *k*-th rectangular spatial window, with (*a*_*k*_, *b*_*k*_) and (*c*_*k*_, *d*_*k*_) specifying its horizontal and vertical bounds, respectively.

### A.3 Feature normalization

Before computing distances or training *f*_*θ*_, we normalize all embeddings to ensure that feature scales are comparable across domains. We standardize features using a target-informed scaler: we compute the mean and standard deviation of each feature over *χ*^*t*^, transform both*χ* ^*s*^ and *χ* ^*t*^ into this standardized space, and apply *f*_*θ*_ in the standardized coordinates. After transformation, we invert the standardization to return *f*_*θ*_( *χ*^*s*^) to the original embedding units. This procedure stabilizes optimization and prevents the M-step from exploiting arbitrary scale differences between domains.

### A.4 Hyperparameters and Hardware Set-up

Key hyperparameters and default intuitions are as follows:

- **FGW trade-off** *α*. Controls the balance between feature similarity and structural similarity. Values in the range *α* ≈ 0.3–0.7 work well in practice.
- **Auxiliary vs. feature weight** *γ*. Balances the auxiliary cost *M*_aux_ and feature cost *M*_feat_ in *M* = *γM*_aux_ + (1 − *γ*)*M*_feat_. Larger *γ* places more weight on domain-invariant auxiliary structure (cell types, spatial coordinates), while smaller values emphasize the foundation-model embeddings.
- **Fitting loss weight** λ. Controls the balance between the pointwise alignment term and the variance term in the M-step objective. Larger λ prioritizes making matched source–target pairs close in embedding space, whereas smaller values place more emphasis on matching the overall feature-wise dispersion of *f*_*θ*_( *χ*^*s*^) to that of *χ*^*t*^.

Other optimization settings (entropic regularization in FGW, subsample sizes, learning rate, and number of EM iterations) are fixed to reasonable defaults (sketches of a few thousand points per sample, number of epochs in M-step set to 100 per EM iteration, Adam optimizer with learning rate 10^*−*3^, and 20–25 EM iterations) and kept constant across experiments. Detailed sensitivity analyses for *α, γ*, and λ are provided in Figs. S2, S3, S4, S6.

We implement FGW with GPU-accelerated Sinkhorn iterations and use Adam for the M-step optimization. All experiments use single-precision (FP32), which is sufficient for stable training. For typical embedding dimensions (*d* = 256–1024) and sketch sizes (∼ 5k × 5k), each EM iteration runs under five minutes on a single A6000 GPU.

### A.4 Datasets and Preprocessing

**RNA-seq:** The scRNA-seq and Xenium datasets are obtained from Vannan et al. [15]. We used the scRNA-seq data from the *Vanderbilt* cohort, totaling 49,871 cells. For the Xenium modality, we selected an interstitial lung disease (ILD) sample from the same cohort (VUILD49LA), which contains 38,280 cells profiled for 343 genes and includes matched VisiumHD data for post-integration validation. Both datasets underwent standard scGPT preprocessing to obtain embeddings. To compute cell-phenotype probabilities for *M*_aux_, we trained a logistic-regression classifier on the scRNA-seq expression data to predict 11 broad cell types using normalized expression values of the genes shared between the scRNA-seq and Xenium samples. For subsequent USHER alignment, we mitigate mass imbalance during optimal transport by subsampling the scRNA-seq dataset to match the Xenium marginal cell-type distribution, yielding 26,323 reference cells used for alignment.

**Imaging:** The histopathology images are obtained from Pratapa et al. [23]. Histology was performed on two near-serial sections (10–20 *µ*m apart) that first underwent PhenoCycler-Fusion protein profiling and MALDI-MSI metabolomic profiling. Since H&E obtained post-PCF matched closely with standard H&E image thickness and staining, we use that as the reference. The sections were subsequently H&E stained and scanned at 40× magnification, producing ∼10,000 × 10,000 pixel RGB images. The two images were co-registered using a simple affine transform. We extracted patch-level features using the H-OptimUS-1 vision transformer, which produces a single 1,536-dimensional embedding for each 14 × 14 pixel patch, yielding an embedding grid of approximately 700×700×1536. To reduce spatial resolution while retaining local structure, we aggregated embeddings within non-overlapping 4 × 4 patch bins, resulting in 38,400 aggregated embeddings, each represented by a 1,536-dimensional vector used for USHER alignment. For *M*_aux_, we used the 2D spatial coordinates of the aggregated patches, leveraging the image registration to provide a stable spatial prior for learning *f*_*θ*_ using USHER.

### A.6 scGPT Embeddings on Tangram-Predicted Spatial Expression

To assess whether imputing spatial gene expression with Tangram improves cross-assay comparability for foundation-model embeddings, we generated scGPT representations using Tangram-predicted Xenium expression values and compared them to embeddings computed from the reference scRNA-seq dataset. Xenium data were first aligned to the reference using Tangram, which outputs per-cell predictions of the full RNA expression profile in the scRNA-seq gene space. These predicted gene expression matrices were then passed to scGPT using the same preprocessing and embedding pipeline applied to the real scRNA-seq data.

The resulting scGPT embeddings from Tangram-predicted Xenium profiles did not recover the manifold structure or cell-type organization present in the reference embeddings (Fig. S1(a)). Instead, predicted-expression embeddings occupied distinct regions of scGPT space, indicating a distributional mismatch between Tangram outputs and real scRNA-seq measurements. Integration metrics such as BRAS and iLISI confirmed this observation: Tangram-predicted embeddings showed mixing scores comparable to, or slightly worse than, embeddings computed directly from raw Xenium expression (Fig. S1(b)), suggesting that Tangram imputation does not improve assay alignment for foundation-model embeddings.

To investigate the source of the mismatch, we compared the marginal distributions of real and predicted expression values (Fig. S1c)). Histograms revealed clear shifts in sparsity, dynamic range, and overall scale between reference scRNA-seq and Tangram-predicted profiles. These altered distributions likely fall outside scGPT’s training manifold and thus produce degraded embedding geometry. Collectively, these analyses show that Tangram’s gene-level predictions, while effective for spatial mapping, do not yield expression distributions suitable for scGPT, and consequently fail to improve cross-assay embedding alignment.

---

**Fig. S1:**
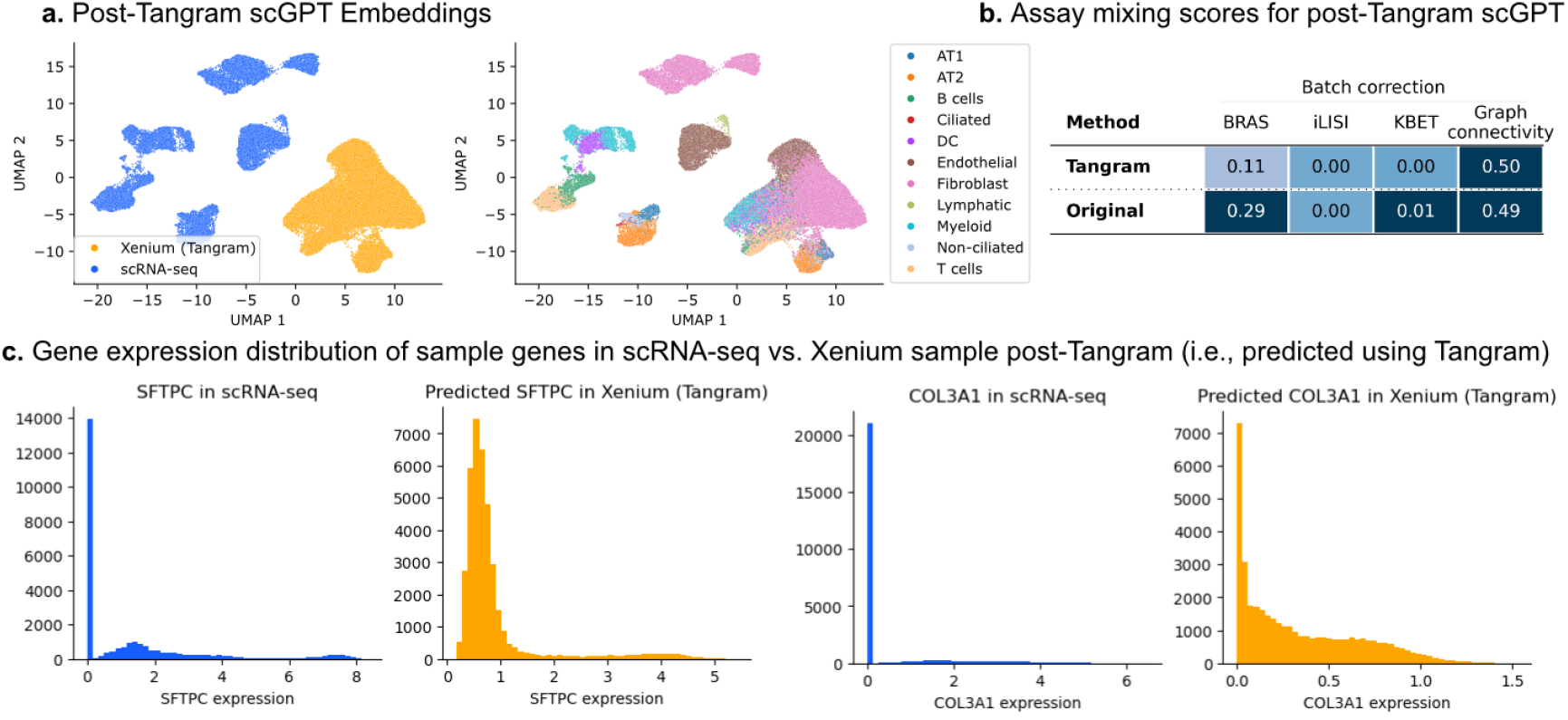
scGPT embeddings computed on Tangram-predicted spatial expression values. **a**. scGPT embeddings computed on Xenium cells post-Tangram alignment fail to produce comparable embeddings to scRNA-seq dataset. **b**. Assay mixing metrics show comparable or degraded performance compared to scGPT embeddings computed on raw expression values (i.e., no Tangram-predicted genes). **c**. Histogram of gene expression values show differences in distributions in reference scRNA-seq dataset vs. Tangram predicted gene expression values.

**Fig. S2:**
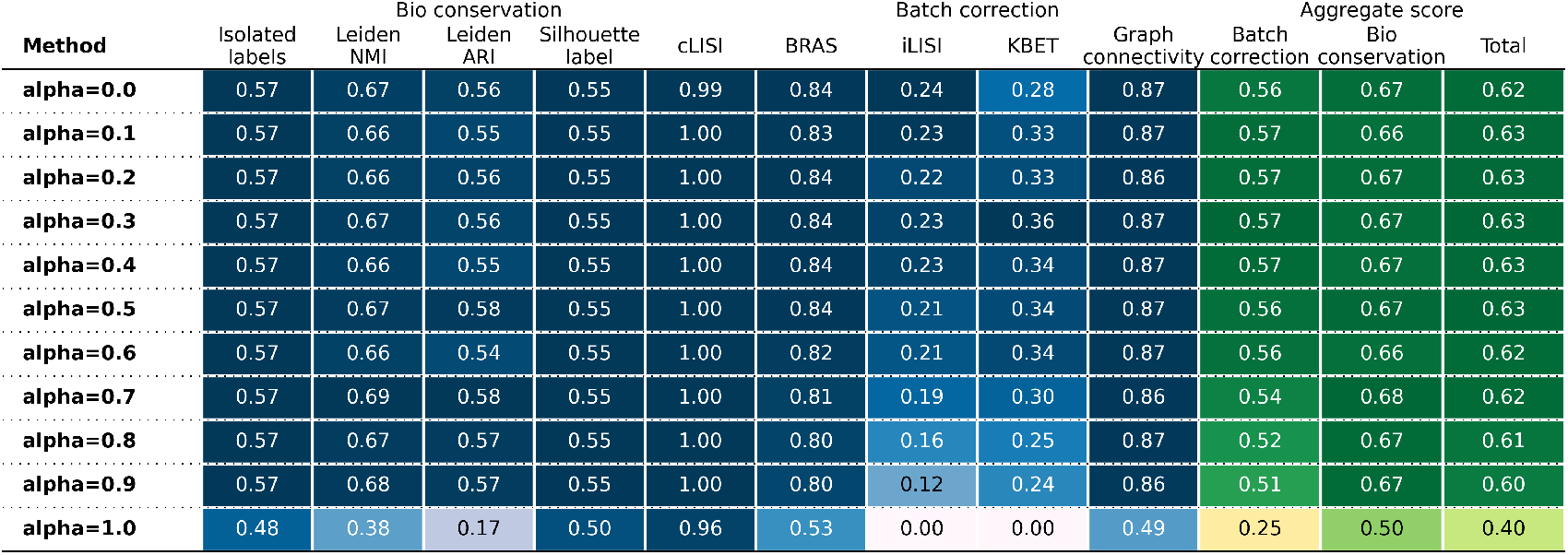
USHER is robust to a choice of *α*’s (for fixed λ = 0.9, *γ* = 0.4), with best results in 0.1-0.9 range and best KBET score (higher=better mixing) at *α* = 0.3.

**Fig. S3:**
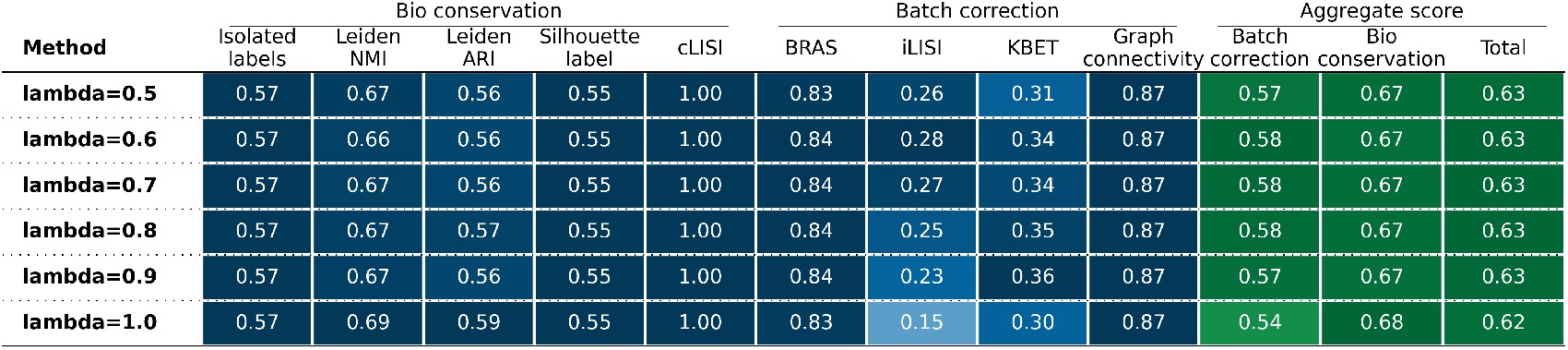
USHER is robust to a choice of λ’s (for fixed *α* = 0.3, *γ* = 0.4), with best batch correction results in a wide 0.6–0.9 range and best KBET score (higher=better mixing) at λ = 0.9.

**Fig. S4:**
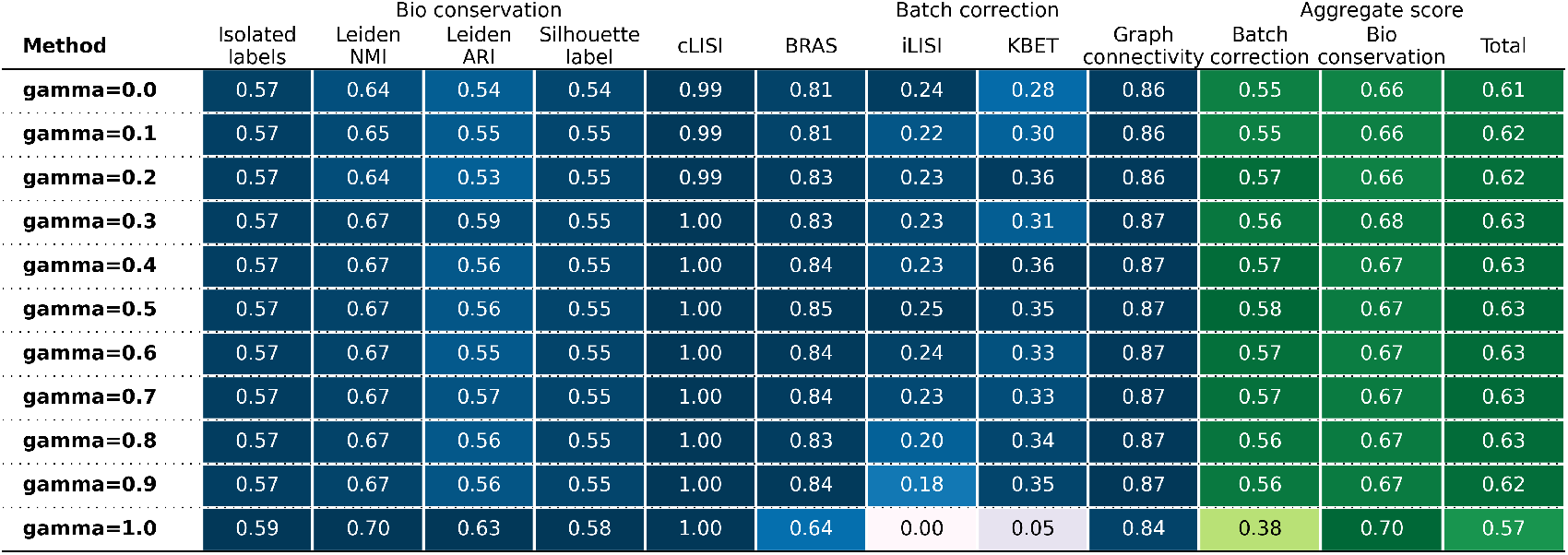
USHER is robust to a choice of *γ*’s (for fixed *α* = 0.3, λ = 0.9), with best batch correction results in a wide 0.0–0.9 range and best BRAS and KBET score (higher=better mixing) at *γ* = 0.4.

**Fig. S5:**
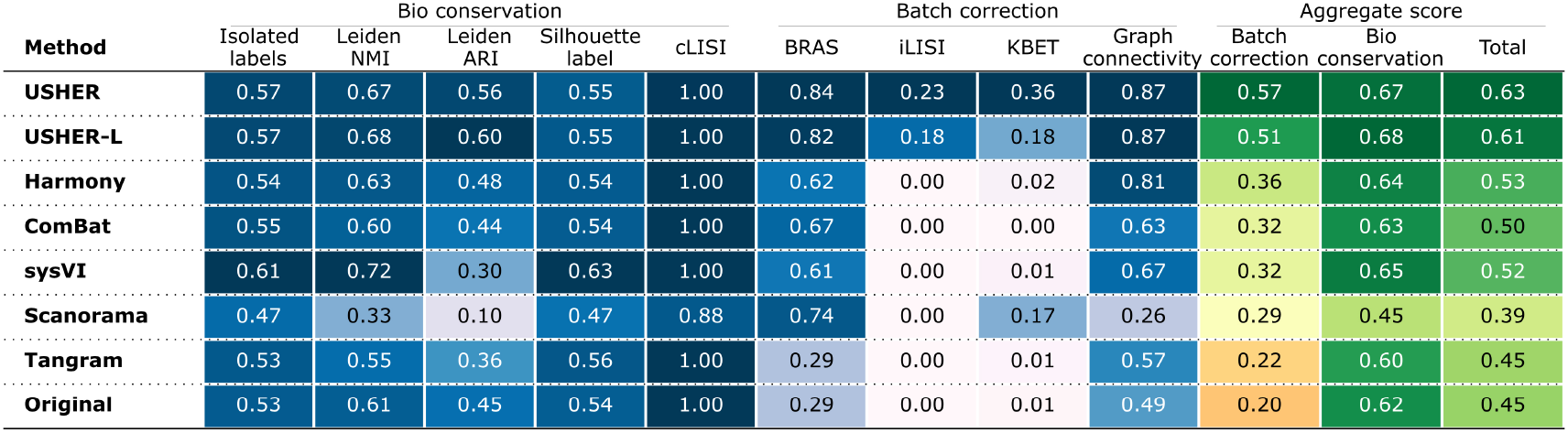
Assay mixing performance of all tested methods, obtained using the ‘scib-metrics’ package.

**Fig. S6:**
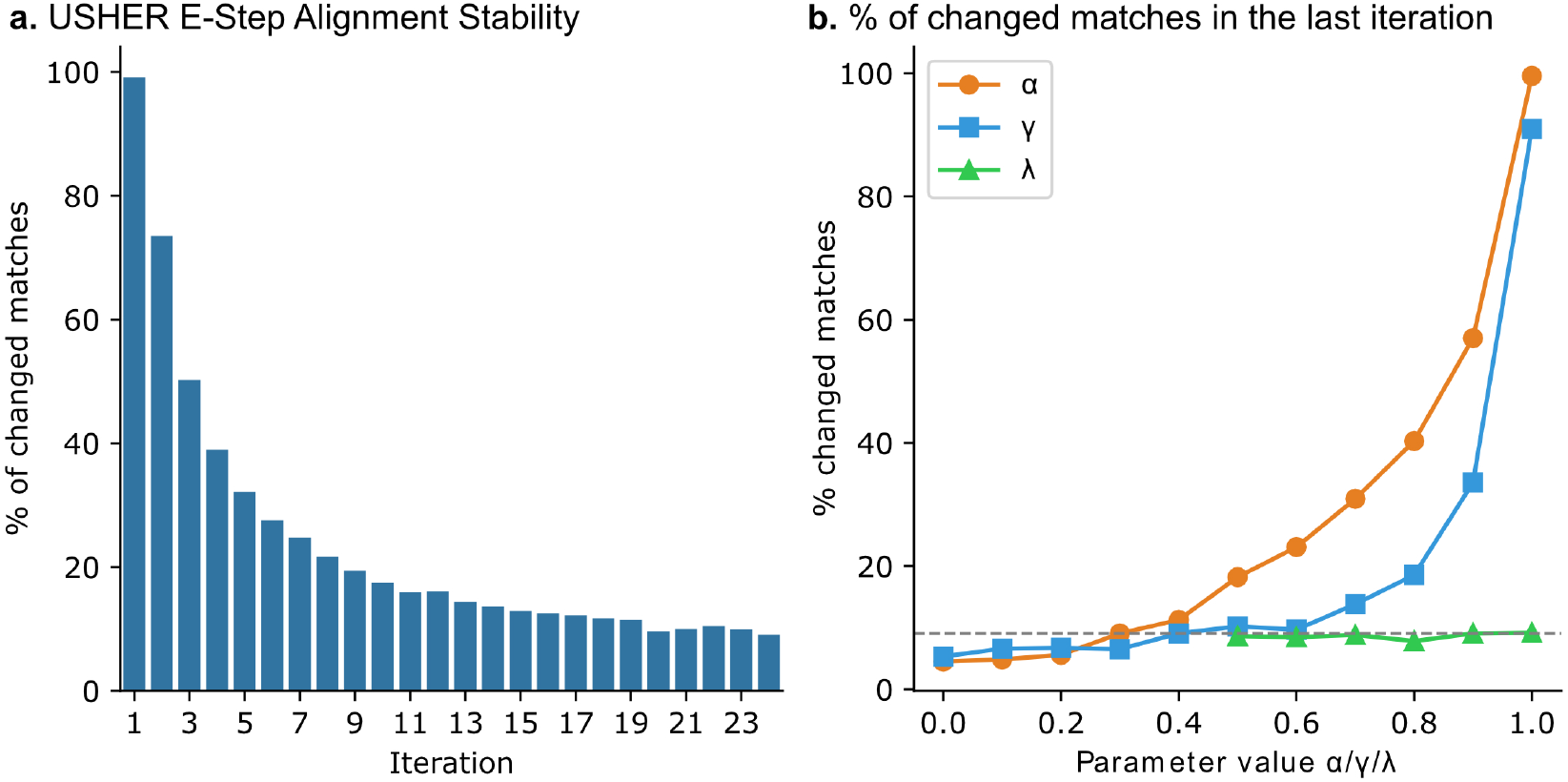
Convergence analysis of USHER’s E-step. **a**. Convergence plot showing the percentage of match changes between successive E-steps under optimal parameters (*α* = 0.3, *γ* = 0.4, λ = 0.9). **b**. Convergence depends strongly on parameter choices. Together, *α* and *γ* control the contribution of the learned representation *f*_*θ*_ to the E-step, with smaller values of either parameter increasing the weight assigned to feature-based matching, resulting in more stable E-step.

**Fig. S7:**
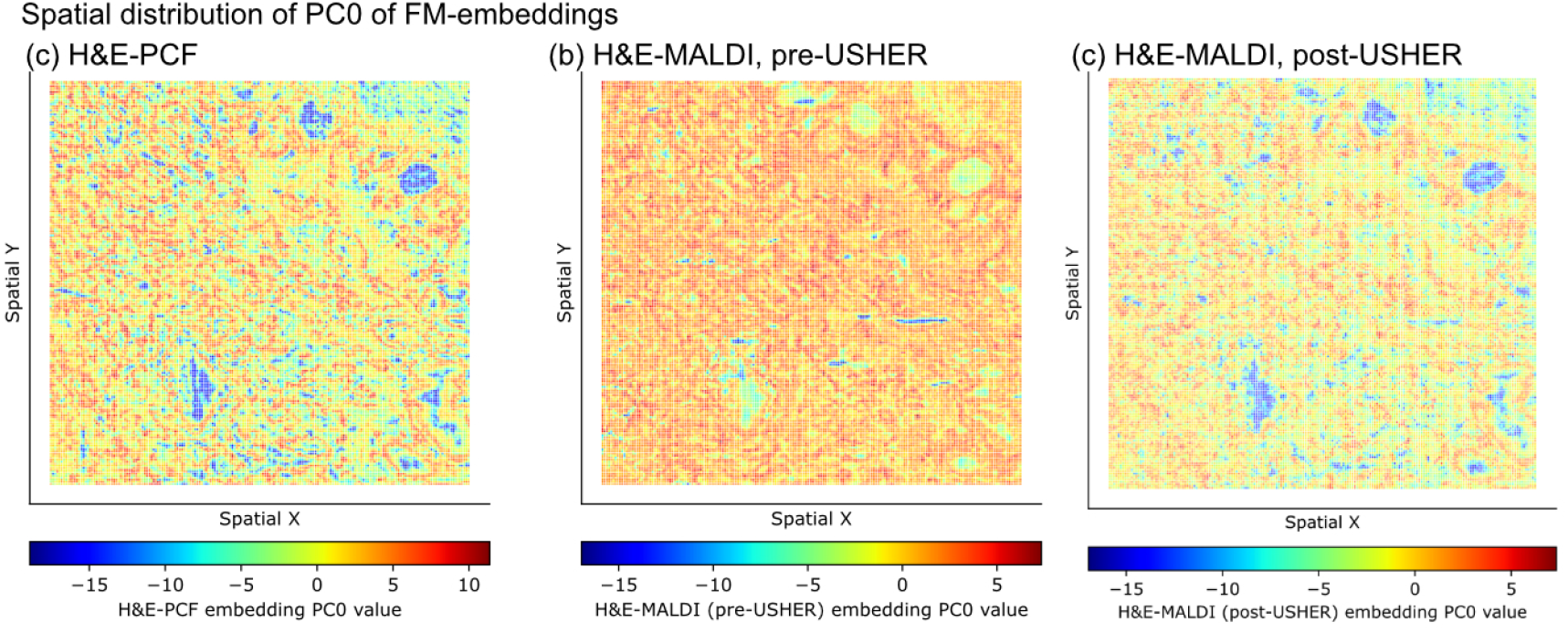
Spatial distribution of principal component PC0 learned from **a**. H&E-PCF FM-embeddings and applied **b**. unmodified pre-USHER H&E-MALDI embeddings, and **c**. post-USHER H&E-MALDI embeddings.

**Fig. S8:**
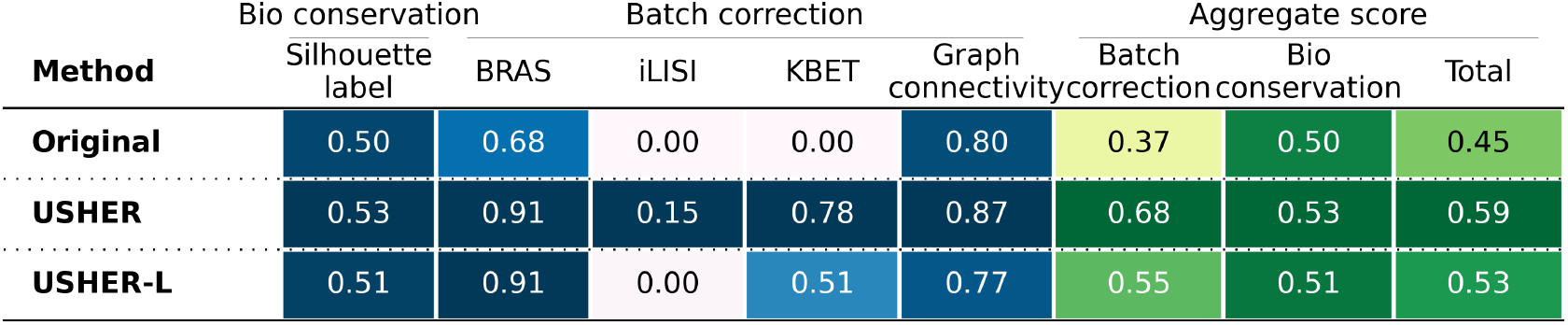
Assay mixing performance of H&E-PCF and H&E-MALDI embeddings before (‘Original’) and after applying USHER (two layer) and USHER-L (single layer), showing improved integration based on different metrics.

